# Rev-Rev Response Element Activity Selection Bias at the HIV Transmission Bottleneck

**DOI:** 10.1101/2023.04.05.535732

**Authors:** Patrick E. H. Jackson, Jordan Holsey, Lauren Turse, Marie-Louise Hammarskjold, David Rekosh

**Author notes:** Corresponding author: Patrick E. H. Jackson 345 Crispell Drive MR6 2524 Charlottesville, Virginia, USA 22908 1-434-982-3559. **Conflict of Interest Statement:** The authors have declared that no conflict of interest exists.

## Abstract

HIV is not efficiently transmitted between hosts, and selection of viral variants occurs during the process of sexual transmission. The factors that confer selective advantage at the transmission bottleneck remain incompletely understood. We explored whether differences in the Rev-Rev Response Element (RRE) regulatory axis of HIV affect transmission fitness, since functional variation in the Rev-RRE axis in different viral isolates has been shown to affect replication kinetics and relative expression of many HIV proteins. Single genome HIV sequences were identified from nine linked subject pairs near the time of female-to-male transmission. Using a rapid flow-cytometric assay, we found that the functional Rev-RRE activity varied significantly between isolates. Moreover, it was generally lower in recipients’ viruses compared to the corresponding donor viruses. In six of nine transmission events, recipient virus Rev-RRE activity clustered at the extreme low end of the range of donor virus activity. Rev-RRE pair activity was an unpredictable product of component Rev and RRE activity variation. These data indicate selection pressure on the Rev-RRE axis during female-to-male sexual transmission. Variation in the activity of the Rev-RRE axis may permit viral adaptation to different fitness landscapes and could play an important role in HIV pathogenesis.

## Introduction

Sexual transmission of HIV is a frequent occurrence worldwide, accounting for more than one million incident infections in 2021 (1). Paradoxically, the virus is not easily transmitted between hosts. The vast majority of unprotected sexual encounters between sero-discordant partners do not result in a new infection (2). Even in cases where sexual transmission does occur, of the many viral variants circulating in the donor partner usually only a single virus successfully enters the new host and establishes infection (3, 4). This sharp reduction in viral diversity between the previously infected donor and the newly infected recipient is termed the transmission bottleneck.

A virus must traverse multiple potential barriers to successfully transmit HIV from a female to a male host via penile-vaginal intercourse (5). The viral quasi-species in the donor’s genital compartment may differ from that of the systemic circulation due to the immune microenvironment, and a subset of genital tissue resident variants may predominate in genital fluids (6). The recipient’s genital mucosa presents a physical obstacle to infection and may also represent an immunologic landscape that differs from that of the donor’s genital immune milieu. The virus must then successfully infect a susceptible cell within the host’s genital tissue and establish productive infection in a regional lymph node. Only then can systemic dissemination occur. Each of these steps – replication in donor genital tissue, entry into donor transmission fluid, infection of a recipient cell past the mucosa, and establishment of productive infection in recipient lymphoid tissue – may or may not individually represent a significant obstacle to transmission. However, the net effect of the transmission bottleneck exerts selection pressure on transmitted/founder (T/F) variants (7).

The T/F virus is generally not the most predominant variant in the donor genital compartment, as would be anticipated if transmission were a mere stochastic process (8). The selected phenotype of the T/F variant is incompletely understood. Selection for CCR5-tropism has been consistently demonstrated across studies (3, 9, 10). Several studies have also observed selection for interferon resistance of T/F variants (11–14), though this remains controversial (15, 16). Phenotypic differences in characteristics such as infectivity (12), virus particle release (11) and envelope content (12) have also been seen in some but not all transmission studies (15, 16). These conflicting results may be due to differences in assay methodology and subject populations. Selection pressures may differ by route of HIV transmission, and factors such as concurrent genital inflammation and high donor viral load may decrease selection stringency (7).

In order to replicate, HIV must overcome the cellular restrictions to the export of intron-containing viral mRNAs from the nucleus to the cytoplasm (17). This is accomplished by means of a *trans*-acting viral protein, Rev, in conjunction with a *cis*-acting RNA secondary structure, the Rev-Response Element (RRE), found in all the viral mRNAs with retained introns (18, 19). Rev is constitutively expressed from a completely spliced viral mRNA. After translation in the cytoplasm, the Rev protein is imported into the nucleus where it binds to the RRE and oligomerizes (20–22). The RRE-Rev complex then recruits cellular factors, including Crm1 and Ran-GTP, to form a complex capable of exporting the intron-containing viral mRNAs to the cytoplasm for translation or packaging into new viral particles (23).

HIV primary isolates exhibit sequence differences in both *rev* and the RRE, and this in turn results in substantial functional activity variation in the Rev-RRE axis between variants (24, 25). Small sequence differences in Rev, the RRE, or both can yield significant differences in Rev-RRE axis activity (26, 27). As the Rev-RRE interaction is necessary for the nuclear export and translation of intron-containing but not fully spliced viral mRNAs, differences in the level of Rev-RRE activity affect not only viral replication kinetics but also the relative expression of many viral proteins (28). Work on another complex retrovirus with a functionally homologous Rev-RRE system, equine infectious anemia virus, has shown that Rev-RRE activity can vary during the course of chronic infection and that the level of activity correlates with clinical disease state (29, 30). Some small studies have also proposed that differences in HIV Rev or RRE activity may affect clinical progression (28, 31, 32). Thus Rev-RRE activity could be a potential factor that contributes to the phenotype of the T/F virus, but this has not been examined to date.

In this study, we examined whether functional differences in the Rev-RRE regulatory axis impact variant fitness at the transmission bottleneck. This was achieved by measuring differences in Rev-RRE functional activity among primary isolates obtained from subjects in linked female-to-male HIV transmission pairs.

## Results

### Identification of Rev-RRE sequences

We selected HIV-1 single genome sequences from eighteen subjects in nine linked female-to-male HIV transmission pairs (Table 1). Eight subjects (four transmission pairs) were participants in the Center for HIV/AIDS Vaccine Immunology (CHAVI-001) acute infection cohort (11, 33) and ten subjects (five transmission pairs) were participants in the Zambia-Emory HIV Research Project (16). All sequences were previously published in GenBank (https://www.ncbi.nlm.nih.gov/genbank/). Each HIV transmission pair was given a code A through I. Recipient samples were obtained during acute infection (Fiebig stage 4 or earlier). The time elapsed between the acquisition of the donor and recipient samples was a maximum of 265 days (median 19 days). All subjects were infected with subtype C virus.

**Table 1.** Sources of HIV genome sequences.

| Pair | Partner | Days between donor and recipient samples | No. single HIV genomes | No. unique Rev aa sequences | No. unique RRE na sequences | No. unique Rev-RRE pairs | Sequence accession numbers | Subject code in reference | Reference |
| --- | --- | --- | --- | --- | --- | --- | --- | --- | --- |
| A | Donor | 19 | 20 | 9 | 12 | 17 | KY112094-KY112126 | CH0212 | Iyer 2017 |
|  | Recipient |  | 60 | 8 | 7 | 14 | JX972986-JX972998,<br>JX973019-JX973074 | CH0162 |  |
| B | Donor | 6 | 18 | 9 | 16 | 17 | KR820367-KR820384 | Z3678F | Deymier 2015 |
|  | Recipient |  | 9 | 1 | 2 | 2 | KR820385-KR820393 | Z3678M |  |
| C | Donor | 47 | 38 | 15 | 16 | 23 | KY112322-KY112359 | CH0492 | Iyer 2017 |
|  | Recipient |  | 33 | 2 | 5 | 6 | KY112218-KY112250 | CH0427 |  |
| D | Donor | 265 | 41 | 6 | 9 | 14 | KY112015-KY112055 | CH1064 | Iyer 2017 |
|  | Recipient |  | 12 | 2 | 3 | 4 | KX216883-KX216893,<br>KX216895 | CH0848 |  |
| E | Donor | 0 | 18 | 8 | 4 | 12 | KR820422-KR820439 | Z4473F | Deymier 2015 |
|  | Recipient |  | 10 | 1 | 2 | 2 | KR820440-KR820449 | Z4473M |  |
| F | Donor | 23 | 17 | 8 | 8 | 13 | KR820341-KR820357 | Z3618F | Deymier 2015 |
|  | Recipient |  | 9 | 2 | 2 | 3 | KR820358-KR820366 | Z3618M |  |
| G | Donor | 65 | 39 | 13 | 27 | 33 | KY112390-KY112428 | CH0596 | Iyer 2017 |
|  | Recipient |  | 18 | 3 | 2 | 4 | KY111965-KY111982 | CH0455 |  |
| H | Donor | 7 | 21 | 8 | 14 | 16 | KR820394-KR820414 | Z4248F | Deymier 2015 |
|  | Recipient |  | 7 | 1 | 1 | 1 | KR820415-KR820421 | Z4248M |  |
| I | Donor | 3 | 20 | 7 | 11 | 15 | KR820294-KR820313 | Z331F | Deymier 2015 |
|  | Recipient |  | 10 | 2 | 1 | 2 | KR820314-KR820323 | Z331M |  |

A total of 400 single genome sequences from these subjects that included the full coding region of *rev* as well as the RRE were analyzed. Phylogenetic trees were generated using the neighbor joining method to confirm the pattern of HIV transmission (Figure 1). Within each transmission pair, sequences from the recipient clustered together and mostly separately from donor sequences. As expected, sequences from different transmission pairs were not interspersed (Figure S1).

**Figure 1.**
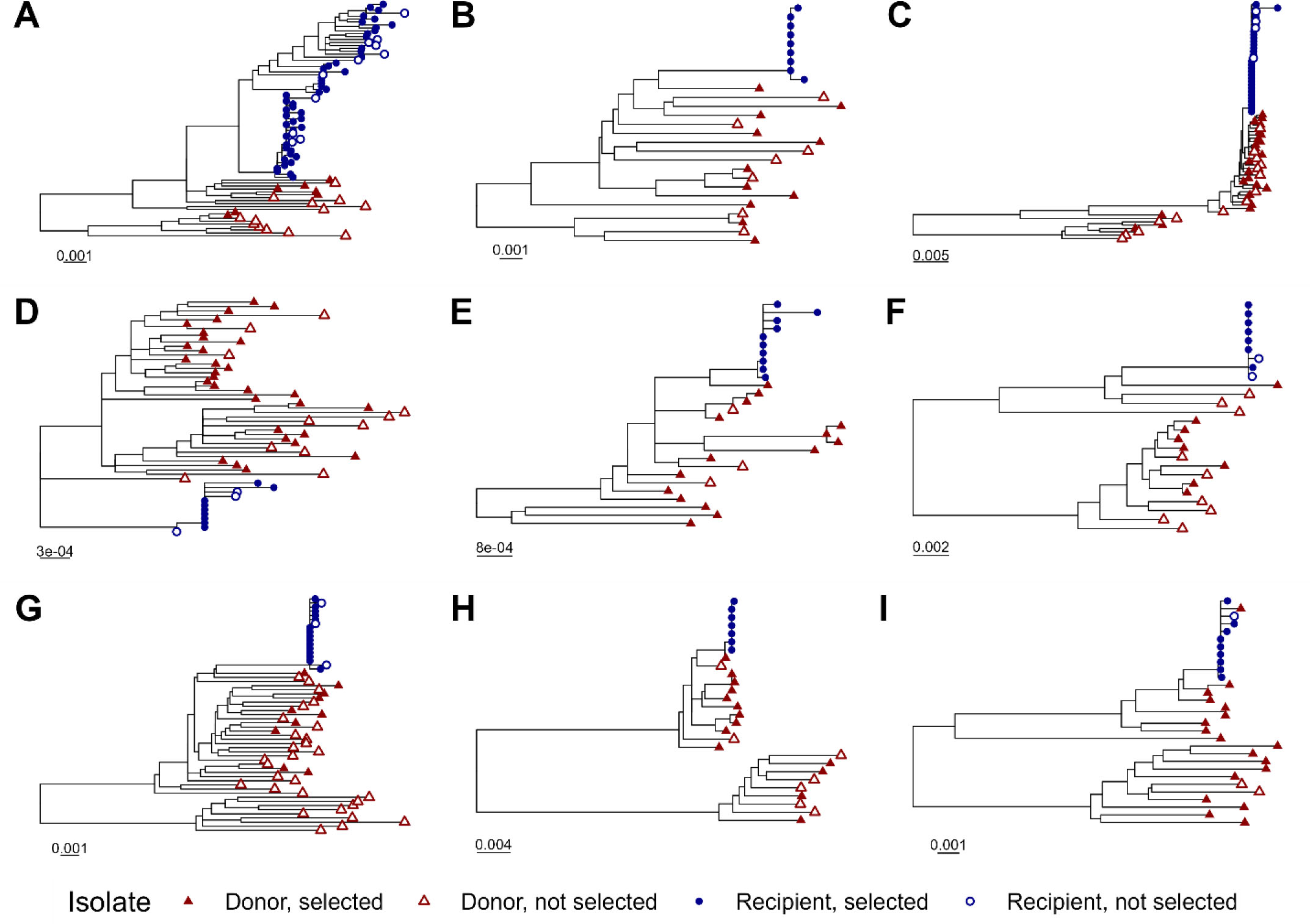
Phylogenetic trees for individual transmission pairs. A phylogenetic tree was generated using the neighbor joining method for 401 single genome HIV sequences. The sequences of four hundred primary isolates associated with eighteen individual subjects in nine linked female-to-male HIV transmission pairs were obtained from GenBank. The laboratory strain NL4-3 was included in tree generation as an outgroup but was excluded from the figure display for clarity. Portions of the tree corresponding to the individual transmission pairs, A through I, are displayed separately. Tip symbols differentiate sequences from donors and recipients, as well as genomes containing Rev-RRE pairs that were selected or not selected for inclusion in functional assays. Horizontal bars represent nucleotide substitutions per site. See also Figure S1.

The 400 analyzed viral genomic sequences included 105 unique Rev amino acid sequences and 142 unique RRE nucleotide sequences. Some RRE and Rev sequences were observed to occur in multiple primary isolates and in multiple combinations. 198 unique Rev-RRE cognate pairs (that is, a Rev amino acid sequence and a RRE nucleotide sequence in a single viral genome) were identified. Donor quasispecies exhibited more sequence diversity than recipient quasispecies. Donor quasispecies had a median of 16 unique Rev-RRE cognate pairs while recipient quasispecies had a median of three unique cognate pairs.

From the set of 198 unique Rev-RRE cognate pairs, a subset was selected for functional analysis based on prevalence within the subject quasispecies. All Rev-RRE cognate pairs present in at least 12% of viral variants circulating in a single host were included in functional assays. Additional Rev-RRE pairs were included to ensure the highest prevalence Rev and RRE sequences in a subject were represented in functional assays. All in all, a total of 81 unique Rev-RRE cognate pairs were utilized in the functional activity assays. Of the original 400 analyzed primary isolates, 281 primary isolates contained a Rev-RRE cognate pair that was represented in the functional assays. One Rev-RRE cognate pair occurred in both the donor and recipient in transmission pair E. All other Rev-RRE cognate pairs occurred in only a single individual.

Between 1 and 13 unique Rev-RRE cognate pairs were selected for each subject, representing coverage varying between 26% and 100% of the sequenced quasispecies in each individual (Figure 2, Figures S2 and S3). For individuals where a few Rev-RRE cognate pairs occurred many times in the quasispecies, a greater degree of coverage was accomplished than when many cognate pairs were present at a low frequency (Table S1). For the donor individuals in transmission pairs A, F, and G, a minority of the circulating variants were represented in the functional assays but all Rev-RRE cognate pairs appearing in the quasispecies more than once were included.

**Figure 2.**
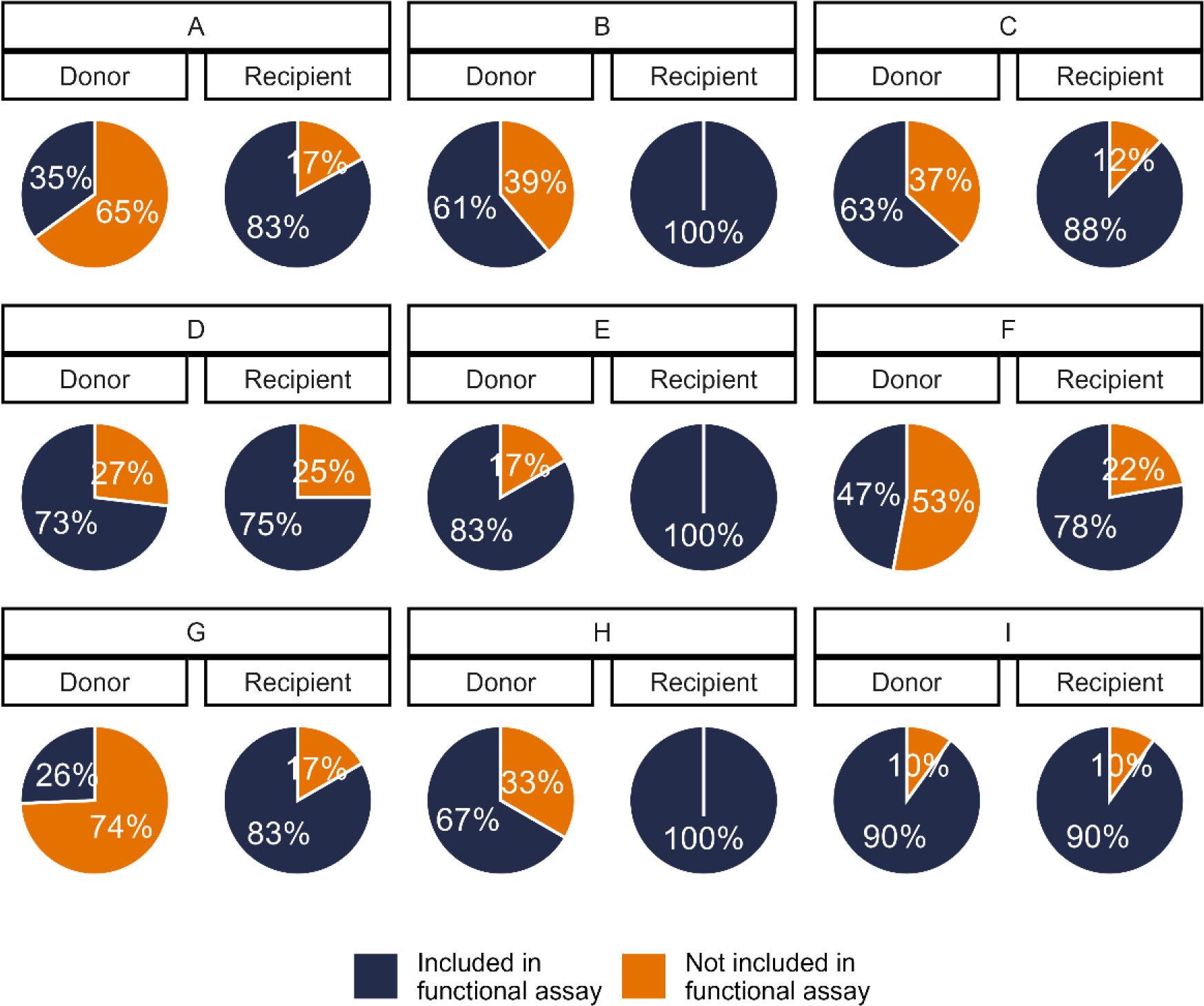
Proportion of viral variants represented in functional activity assays. A subset of the unique Rev-RRE cognate pairs from primary isolates were included in the functional activity assays. For each subject, the proportion of the sequenced viral variants from that individual’s plasma containing a Rev-RRE sequence included in the functional assays is shown.

### Rev-RRE functional activity of donor and recipient viruses

The relative functional activity of the selected Rev-RRE cognate pairs was determined using a lentiviral vector based assay as previously described (see Methods) (27, 34). As shown in Figure 3, there was an almost 9-fold difference in functional activity between the most-active and least-active Rev-RRE cognate pairs. For each transmission pair where multiple recipient Rev-RRE sequences were assayed, the range of Rev-RRE activity of the donor-derived variants was greater than the range of activity for variants from the corresponding recipient (see also Table S1).

**Figure 3.**
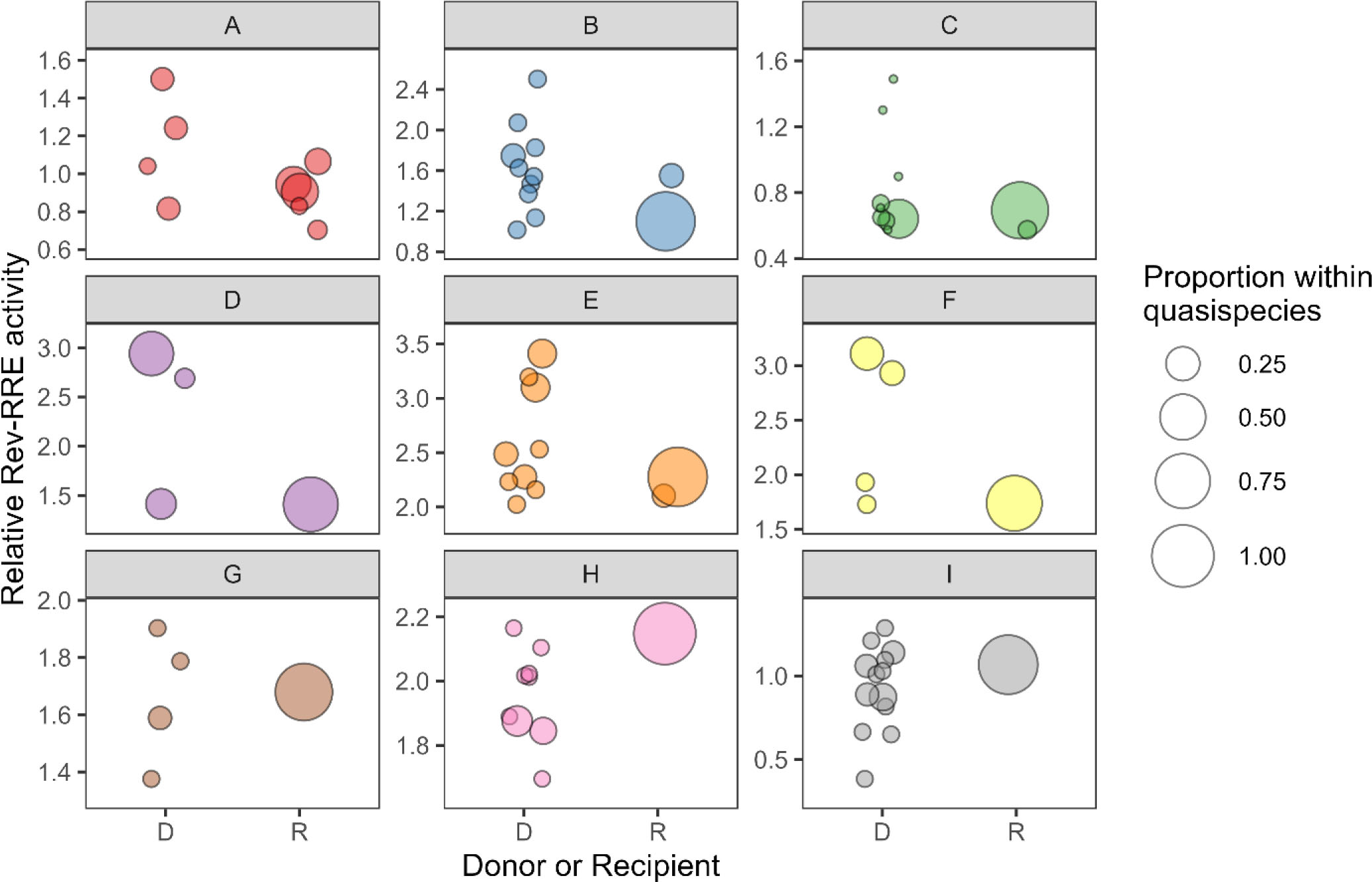
Rev-RRE functional activity of viral variants from donors and recipients. The relative functional activity of selected Rev-RRE pairs from primary isolates in both donors and recipients is shown. Each HIV transmission pair, A through I, is displayed, with Rev-RRE cognate pairs from donor viral sequences on the left of each box and Rev-RRE cognate pairs from recipient viral sequences on the right. Each bubble is a unique Rev-RRE pair in the indicated individual’s quasispecies. The position of the bubble on the y-axis represents the relative level of Rev-RRE functional activity for that pair. The area of each bubble is scaled to the relative prevalence of the Rev-RRE pair sequence within the individual’s sequenced quasispecies. Activity units are multiples of the Rev-RRE activity of the laboratory strain NL4-3. D – donor, R – recipient.

Overall, the Rev-RRE activity of recipient-derived variants was significantly lower than the activity of the corresponding donor variants (*p* = 0.02). In six of the nine transmission pairs, the Rev-RRE activity of the recipient variants clustered at the extreme low end of the range of activity of the corresponding donor variants. This pattern was consistent for transmission pairs with overall high Rev-RRE activity (pairs D, E, F, G, and H) and transmission pairs with overall low Rev-RRE activity (pairs A, B, C, and I). There was no transmission pair where a recipient variant had the highest overall Rev-RRE activity.

In only three of the nine transmission pairs - G, H, and I – was the weighted average of recipient variant Rev-RRE activity greater than the average donor variant activity. It should be noted that for one of these transmission pairs, I, the variant Rev-RRE sequences tested had the lowest overall level of activity of any transmission pair (Figure S4).

### Rev and RRE contributions to cognate pair activity

To evaluate the contribution of Rev and RRE differences to cognate pair activity, artificial combinations of Revs and RREs were next tested in the functional assay. Artificial Rev-RRE pairs consisted of a Rev sequence from the HIV laboratory strain NL4-3 paired with an RRE from a primary isolate, or a Rev from a primary isolate paired with the NL4-3 RRE. For five primary isolate Rev-RRE cognate pairs, functional assays were performed comparing the activity of the primary isolate cognate pairs with these artificial pairs (Figure 4).

**Figure 4.**
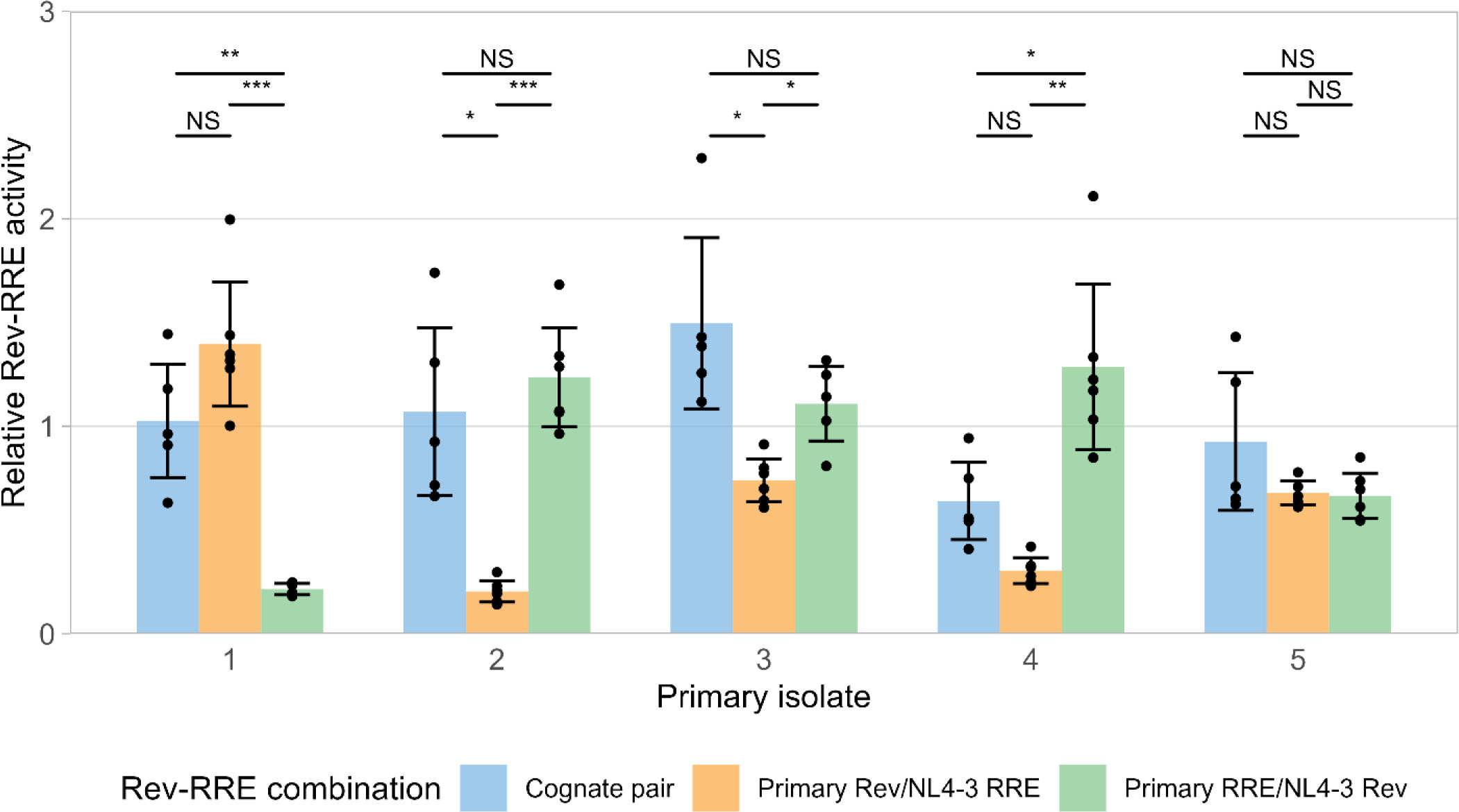
Contributions of Rev and the RRE to cognate pair functional activity. The relative functional activity of Rev-RRE cognate pairs from primary isolates, as well as the activity of the component Rev with NL4-3 RRE and the component RRE with NL4-3 Rev is shown. Relative activity is shown as multiples of the activity of the NL4-3 Rev/NL4-3 RRE cognate pair without units. Observations from technical replicates (n = 5 or 6) are shown as dots. Bars represent the mean value of individual observations and error bars represent standard deviation. Statistical comparisons were performed using a one-way ANOVA with adjustment for multiple comparisons using Dunnett’s T3 method. Primary isolate 1 was obtained from donor I (accession KR820312), isolate 2 was obtained from recipient B (KR820385), isolate 3 was obtained from donor G (KY112428), isolate 4 was obtained from donor C (KY112275), and isolate 5 was obtained from recipient A (JX973051). Cognate pair activity values replicate values shown in Figure 3. NS not significant, * p <0.05, ** p <0.01, *** p<0.001.

Cognate pair functional activity could not be predicted by component Rev or RRE activity. For example, Rev-RRE cognate pair 1 exhibited high Rev activity and low RRE activity, while cognate pair 2 displayed low Rev and high RRE activity. This is consistent with previous results indicating that changes in either Rev or the RRE are sufficient to significantly alter Rev-RRE axis activity.

## Discussion

In this study, we observed a previously unreported selection based on the Rev-RRE axis during female to male sexual transmission of HIV. Globally, recipient viral variants displayed lower Rev-RRE activity than variants from corresponding donors, and in six of nine transmission events recipient variant activity clustered at the extreme low end of the range of activity of the corresponding donor variants. Recipient variant Rev-RRE activity was generally not the same as the activity of the predominant variant in the donor plasma compartment. These results are most consistent with the female-to-male sexual transmission bottleneck conferring a selection advantage for viruses with a lower level of Rev-RRE activity.

There were three discordant observations, however, where the weighted average of recipient variant Rev-RRE activity was higher than that of the corresponding donor. For transmission pair I, the Rev-RRE activity of variants from both the donor and recipient were low compared with the set of variants from all transmission pairs. If Rev-RRE selection at the transmission bottleneck is subject to a threshold effect, then the activity of these donor viruses may have been sufficiently low that there was no additional selection during transmission. While this potential explanation would not account for transmission pairs G and H, in which variants displayed intermediate levels of Rev-RRE activity, selection pressures at the transmission bottleneck may be mitigated by conditions that predispose to forward transmission including concurrent genital inflammation (7) and immune compromise. In the absence of additional clinical data for these subjects, it is unclear if these factors could account for the cases where recipient Rev-RRE activity was higher.

The pattern of donor and recipient virus Rev-RRE activity is unlikely to be secondary to selection for other phenotypic parameters such as coreceptor utilization or interferon resistance. As demonstrated previously and again in this study, Rev-RRE activity is highly sensitive to changes in the RRE, *rev*, or both. This permits a high degree of plasticity in the regulatory axis and an ability to accommodate extrinsic pressures while maintaining Rev-RRE activity level. Previous work by Frankel *et al.* has demonstrated that functional regions of *rev* are segregated from functional regions of the overlying *tat* and *env* genes, permitting mutations that affect the activity of one protein at a time (35, 36). RRE activity is similarly robust to nonsynonymous changes in Env, as the functional consequence of these mutations can be compensated by additional synonymous changes in other portions of the RRE (37, 38). Therefore, a constraint on Env sequence imposed by a selection for type I interferon resistance at the transmission bottleneck, for example, would not be expected to impose a constraint on T/F Rev-RRE activity level. Indeed, Iyer *et al.* experimentally demonstrated widely disparate levels of INFα2 and INFβ sensitivity among these primary isolates, some of which share identical Rev-RRE cognate pairs (11).

An analysis of larger numbers of female-to-male transmission pairs are necessary to validate the finding of apparent selection pressure on the Rev-RRE axis described here. Future studies should ideally also include viral variants of different subtypes, since these may be subject to different selection pressures (12). Other routes of HIV transmission, such as male-to-female sexual transmission, transmission via insertive or receptive anal intercourse, vertical transmission, and parenteral transmission may be subject to differing selection pressures. In this study, we did not assess other viral factors that may contribute to transmission fitness, such as replication capacity, but this may be impacted directly by Rev-RRE activity. A more complete phenotypic characterization of T/F variants may yield insights into the optimal constellation of viral characteristics that facilitate sexual transmission.

In this study, we conceptualized the transmission bottleneck as the sum total effect of multiple potential immunologic and anatomic barriers to the sexual transmission of HIV. We did not assess the contribution of individual potential barriers to selection pressure on the T/F virus. The selection we observe could occur entirely within the donor or the recipient, or selection could be an emergent phenomenon of processes within both hosts. As in other clinical studies of HIV transmission, the single T/F virus was not directly sampled, and viral sequences from the recipient close to the time of transmission were used as a proxy for determining the characteristics of the T/F variant. The limited genetic and functional diversity of recipient viruses suggests that this approach was successful in the present study. However, we cannot exclude rapid selection on the Rev-RRE system occurring during early dissemination in the new host accounting for these findings, rather than selection occurring during the initial sexual transmission.

While we did not examine the mechanism by which Rev-RRE activity selection occurs during transmission, two factors may yield a selective advantage for variants with different levels of Rev-RRE functional activity. First, the Rev-RRE axis impacts viral replication capacity. We previously demonstrated with replication-competent HIV constructs that Rev activity is positively correlated with replication kinetics (27). Second, Rev-RRE activity alters the relative expression of viral proteins encoded by completely spliced mRNA species (*i.e.* Tat, Rev, and Nef) to proteins encoded by mRNAs with retained introns (*e.g.* Gag, Env) (28). Nef modulates the immune response to HIV infection by downregulating CD4 and major histocompatibility complex class 1 expression on the surface of infected cells (39). By maintaining a protective level of Nef expression (independent of Rev-RRE function) and lower expression of the Rev-RRE dependent structural proteins that generate antigenic peptides, viruses with low Rev-RRE functional activity appear to be relatively protected from cytotoxic T-cell-mediated killing (28). In the context of the female-to-male transmission bottleneck, such an immune evasive strategy may explain the selective advantage for variants with lower Rev-RRE activity.

This study sheds light on two areas of investigation. First, we suggest Rev-RRE activity as a new factor in phenotypic selection at the HIV transmission bottleneck. If additional studies confirm this finding, accounting for Rev-RRE activity variation may help to reconcile the currently conflicting data on viral transmission fitness and could potentially point to new strategies for transmission prevention. Second, this study adds to the literature suggesting a role for variation in the Rev-RRE system in HIV pathogenesis (40) and strengthens the concept that the Rev-RRE axis can function as a molecular rheostat to allow the virus to adapt to different pressures. This paradigm may not only be important in HIV transmission but also in other processes where viral adaptation to differing immune environments could affect pathogenesis, including viral compartmentalization and latency.

## Methods

### Sequence selection and processing

Single genome HIV sequences from eighteen individuals consisting of nine female-to-male transmission pairs were identified using the Los Alamos HIV Sequence Database (http://www.hiv.lanl.gov/) and GenBank (41). The sequences were previously published by others) (11, 16) (see Tables 1 and S2 for accession numbers). Only single viral genomes sequenced from plasma that included the RRE and both exons of *rev* were utilized in this study. RRE and *rev* nucleotide sequences were extracted from the original sequence record using the Gene Cutter tool from the Los Alamos HIV database (https://www.hiv.lanl.gov/content/sequence/GENE_CUTTER/cutter.html). Additional sequence analysis and manipulation was performed using Geneious Prime (Dotmatics). Sequences were assessed for clear errors (*i.e.* a premature stop codon at position <100 in *rev,* stop codons in all three forward reading frames within the RRE) and were excluded from further analysis if either of these conditions were met. Rev open reading frames were extracted from the Gene Cutter output.

For each subject, unique Rev amino acid and unique RRE nucleotide sequence pairs were identified within the set of complete genomic sequences. Additionally, unique Rev-RRE cognate pairs (that is, the unique combination of a Rev amino acid sequence and an RRE nucleotide sequence in the same viral genome) were identified. The relative prevalence of unique Revs, RREs, and Rev-RRE pairs within a subject quasispecies was calculated as the number of viral genomes in which this sequence occurred, divided by the total number of viral genomes with intact Rev and RRE sequences in that individual.

All unique Rev-RRE pairs found in at least 12% of circulating variants within an individual quasispecies were included in functional assays. Additional Rev-RRE pairs were selected for functional assays based on Rev or RRE prevalence. No prediction of Rev-RRE functional activity was performed prior to selecting sequences for inclusion in functional assays.

### Phylogenetic analysis

Phylogenetic trees were generated using the viral genomic sequences listed in Table 1. Sequences were aligned in Geneious Prime using the Clustal Omega 1.2.2 algorithm (42). The sequence of the laboratory HIV strain NL4-3 (GenBank accession U26942.1) was included as an outgroup (43). A neighbor-joining phylogenetic tree was generated using the TreeMaker tool from the Los Alamos HIV database (https://www.hiv.lanl.gov/components/sequence/HIV/treemaker/treemaker.html) utilizing a Jukes-Cantor distance model with equal site rate. Tree visualizations were created using R version 4.2.1 and the package ggtree (44) (Figures 1, S1). The NL4-3 tip was removed from tree visualizations for clarity.

### Functional assays

Rev-RRE functional activity assays were performed using a flow cytometry-based system that has been previously described (34). This system includes two packageable vector constructs. The first construct is an NL4-3-derived HIV construct with modifications to render it replication incompetent and to silence native *rev* expression. The construct expresses an eGFP fluorescent marker from the *gag* open reading frame in a Rev-RRE dependent fashion and an mCherry fluorescent marker from the *nef* open reading frame in a Rev-RRE independent fashion. The RRE sequences are flanked by restriction sites for exchange in the native position within *env*. The second assay construct is derived from a murine stem cell virus (MSCV) vector. This construct is modified to express an exchangeable Rev along with an eBFP2 fluorescent marker from a bicistronic construct. The plasmid constructs utilized in these experiments are listed in Table S3.

Selected Rev and RRE sequences were commercially synthesized and cloned into MSCV and HIV vector constructs, respectively. The constructs were then packaged and pseudotyped with VSV-G in 293T/17 cells. To perform the functional assays, SupT1 cells were co-transduced with one Rev- and one RRE-containing assay construct. Transductions were performed in 96 well plates with 2.5 x 10^5^ SupT1 cells in each well. Cells were transduced at a target multiplicity of infection (MOI) of 0.18. Transductions were performed by combining cells, vector stocks, and 8 mcg/mL DEAE-dextran, and then centrifuging the cultures at 380 RCF for 1 hour at room temperature. Flow cytometry was performed 72 hours after transduction using an Attune NxT flow cytometer with autosampler (Thermo Fischer Scientific). Post-acquisition color compensation and data analysis was performed using FlowJo v10.6.1 (FlowJo, LLC).

To analyze flow cytometry data, gates were constructed to define a single cell population. Next, a gate was constructed to include only cells successfully co-transduced with both the Rev- and RRE-containing assay constructs and expressing both mCherry and eBFP2. In this final population, the mean fluorescence intensity (MFI) of eGFP and eBFP2 was determined. Relative Rev-RRE activity was calculated as the ratio of eGFP to eBFP2 MFI for each well.

For each experimental run in which a particular Rev-RRE pair was assayed, three replicate wells were transduced with the same vector constructs. Individual wells were excluded from analysis if more than 32% of cells were positive for either vector construct or if fewer than 500 cells were successfully co-transduced with both constructs. For every experimental run, the mean activity measurement of all interpretable wells for a particular Rev-RRE pair was calculated. Only experimental runs in which at least two wells containing a particular Rev-RRE pair were interpretable were used to contribute data for the activity of that pair. A single experimental run including two or three wells transduced with Rev-RRE pair was considered a single technical replicate for the purposes of statistical analysis.

### Statistics

The relative functional activity of Rev-RRE pairs was consistent across experimental runs, but the absolute value of the eGFP:eBFP2 ratio used as the measurement of activity level varied between experimental runs presumably due to differences in cell line passage. Relative Rev-RRE activity was compared between unique cognate pairs as in Figure 3 using a linear mixed model by restricted maximum likelihood where an experimental run was considered as a random effect and Rev-RRE pair as a fixed effect. Statistical analysis was performed using R version 4.1.2 and the lme4 (45) and lmerTest packages (46). Relative Rev-RRE cognate pair activity was calculated from all available experimental runs, and a minimum of three experimental runs (i.e. three technical replicates) was included for each pair in the model.

To compare Rev-RRE activity between donor and recipient quasispecies, the lme4 package was used to model variant activity with donor vs recipient status as a fixed effect and transmission pair as a random effect. Rev-RRE pair activity values were weighted according to the frequency of occurrence within an individual quasispecies. The estimated marginal means were then calculated and compared for donors versus recipients using R version 4.1.2 and emmeans package (47).

To compare the activity of Rev-RRE cognate pairs and corresponding artificial pairs including either the NL4-3 Rev or the NL4-3 RRE as shown in Figure 4, activity measurements for each Rev-RRE pair were normalized to the activity of the NL4-3 Rev-RRE cognate pair that was included in the same experimental run. Analysis of the difference between cognate pair activity and the activity of the corresponding NL4-3 Rev/primary RRE and primary RRE/NL4-3 Rev pairs was conducted using a one-way ANOVA test with adjustment for multiple comparisons using Dunnett’s T3 method. Statistical analysis was performed using SPSS (IBM).

## Author contributions

P. E. H. J. contributed to conceptualization of the study, performed functional assays, and wrote the first draft of the manuscript. J. H. and L. T. performed functional assays. D.R. and M.-L. H were involved in the conceptualization of the studies and the experimental design, and critically revised the manuscript. All authors reviewed the final manuscript.

## Acknowledgements

The authors acknowledge the University of Virginia Flow Cytometry Core (RRID: SCR_017829) for assistance with performance of functional assays and Clay Ford of the University of Virginia Library for assistance with statistical analysis. Flow cytometry experiments were supported in part by the National Cancer Institute P30-CA044579 Center Grant. Research on HIV Rev was supported by grants CA206275, AI136671 and AI134208 from the National Institutes of Health and the Myles H. Thaler Research Support Gift at the University of Virginia. Partial salary support for P. E. H. J. was provided by grant K08 AI136671 from the National Institutes of Health, for D.R. by the Myles H. Thaler Professorship at the University of Virginia, and for M.-L. H. by the Charles H. Ross Jr. Professorship at the University of Virginia.

## Notes

### Competing Interest Statement

The authors have declared no competing interest.

