## Supplemental figures for "Rev-Rev Response Element Activity Selection Bias at the HIV Transmission Bottleneck"

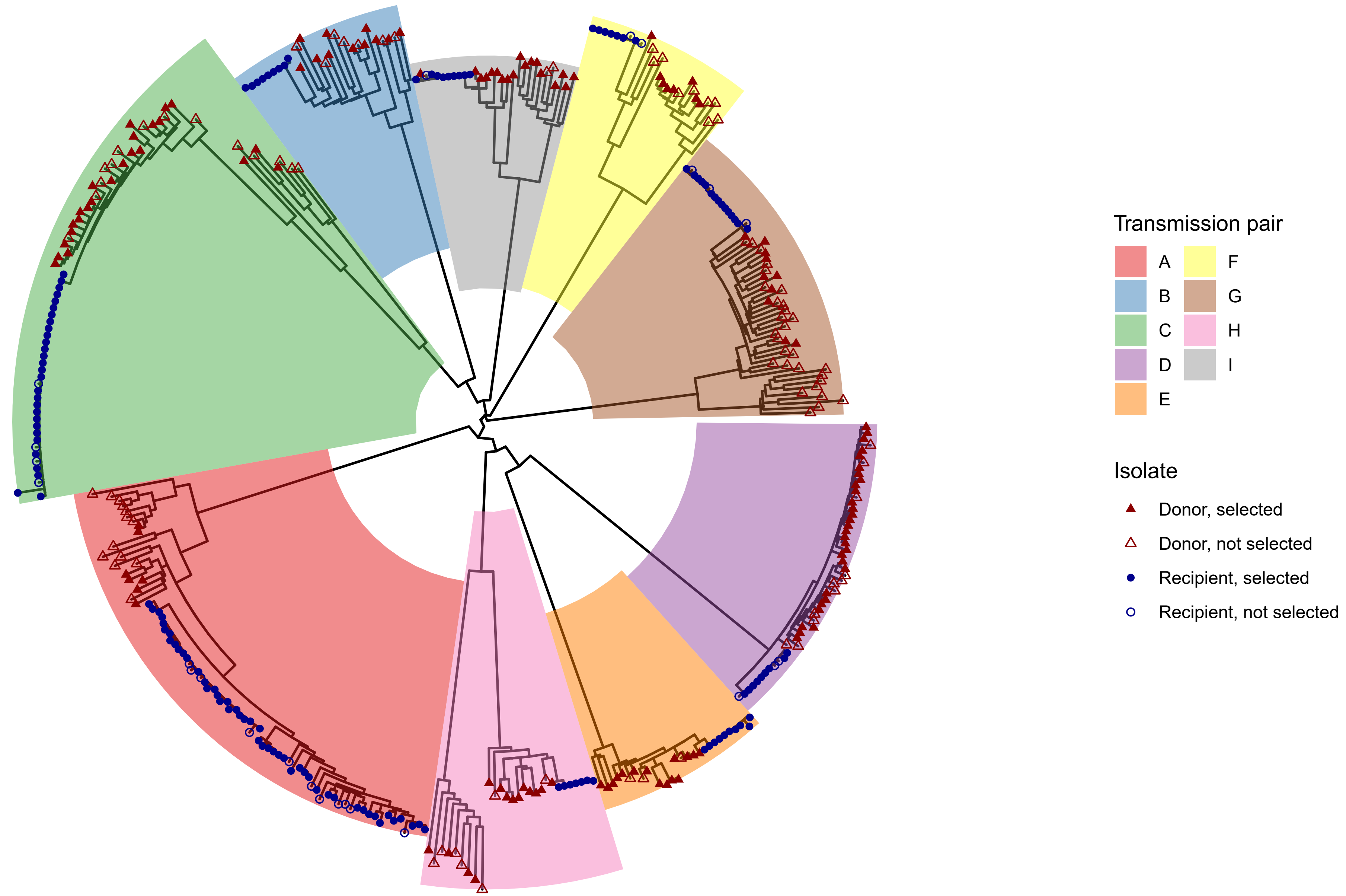


Figure S1. Phylogenetic tree of single genome HIV sequences from eighteen subjects. A phylogenetic tree was generated using the neighbor joining method for 401 single genome HIV sequences. The sequences of four hundred primary isolates associated with eighteen individual subjects in nine linked female-to-male HIV transmission pairs were obtained from GenBank. The laboratory strain NL4-3 was included in tree generation as an outgroup but was excluded from the figure display for clarity. Branches corresponding to each transmission pair, A through I, are differentiated by colored fields. Tip symbols differentiate sequences from donors and recipients, as well as genomes containing Rev-RRE pairs that were selected or not selected for inclusion in functional assays.

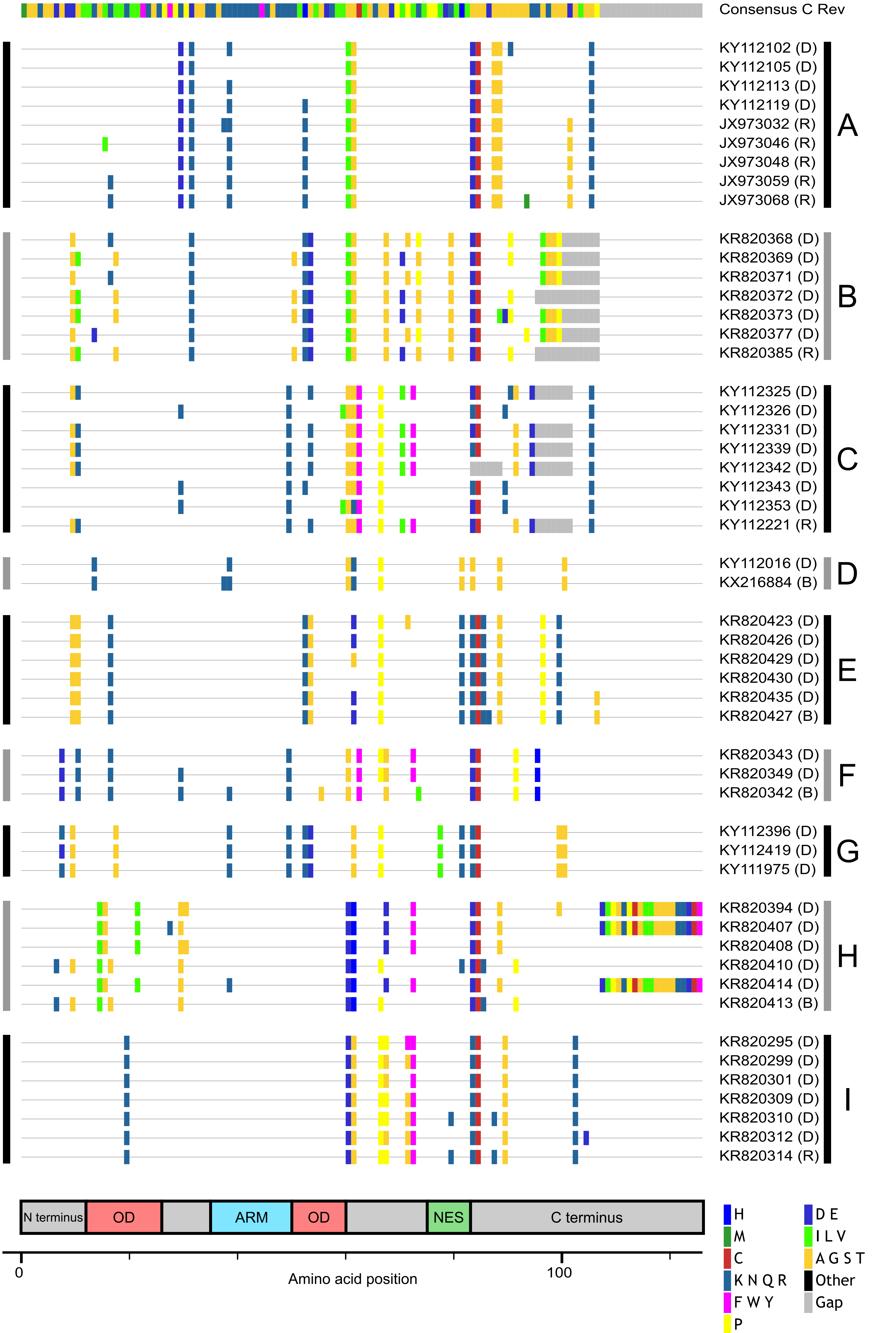


Figure S2. Alignment of unique Rev sequences included in functional assays. Fifty-one unique Rev amino acid sequences from primary isolates that were included in functional assays were aligned along with the Rev from a subtype C consensus sequence (Los Alamos HIV database 2002 consensus, https://www.hiv.lanl.gov/content/sequence/NEWALIGN/align.html). Mismatches between the indicated primary isolate Rev amino acid sequence and the consensus subtype C Rev sequence are shown with vertical bars colored by the residue change or gap. Accession numbers are provided as an example of one primary isolate in which the corresponding Rev sequence is found; other primary isolates may also share the same sequence. Sequences are noted as occurring in only donor (D), only recipient (R), or both donor and recipient (B) subjects within a transmission pair. The transmission pair from which the isolate was sequenced is indicated by labeled vertical black and gray bars to the left and right of the sequences. No Rev sequences occurred in multiple transmission pairs. Amino acid positions corresponding to functional domains of Rev are indicated by the boxes at the bottom of the figure. Functional domains are colored for clarity. OD – oligomerization domain; ARM – arginine rich motif; NES – nuclear export signal.

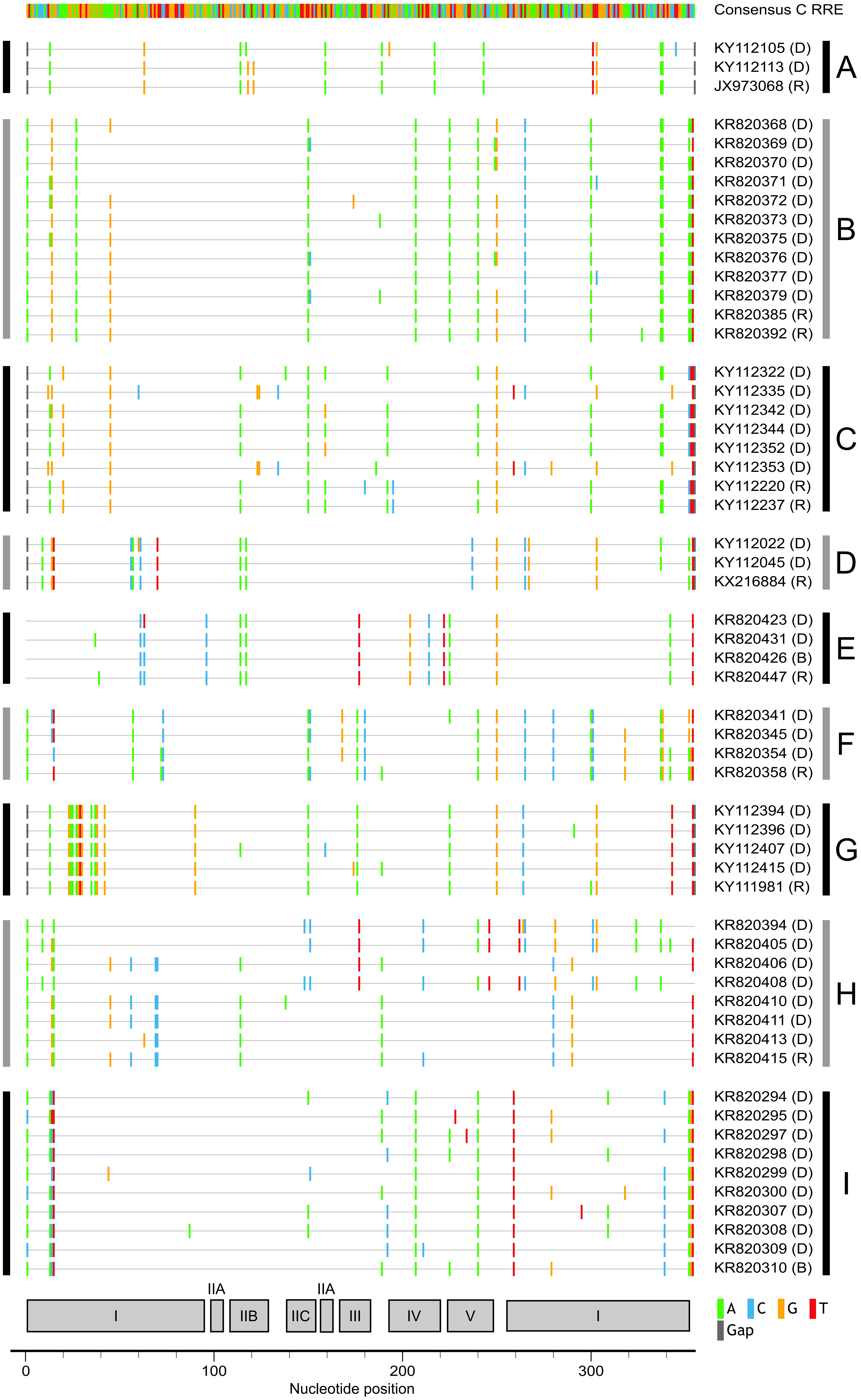


Figure S3. Alignment of unique RRE sequences included in functional assays. Fifty-seven unique RRE nucleotide sequences from primary isolates that were included in functional assays were aligned along with the RRE from a subtype C consensus sequence (Los Alamos HIV database 2002 consensus, https://www.hiv.lanl.gov/content/sequence/NEWALIGN/align.html). Mismatches between the indicated primary isolate RRE sequence and the consensus subtype C RRE sequence are shown with vertical bars colored by the nucleotide change or gap. Accession numbers are provided as an example of one primary isolate in which the corresponding RRE sequence is found; other primary isolates may also share the same sequence. Sequences are noted as occurring in only donor (D), only recipient (R), or both donor and recipient (B) subjects within a transmission pair. The transmission pair from which the isolate was sequenced is indicated by labeled vertical black and gray bars to the left and right of the sequences. No RRE sequences occurred in multiple transmission pairs. Gray boxes at the bottom of the figure indicate nucleotide positions that correspond to the specified stem-loop (I through V) in the NL4-3 five stem-loop structure.

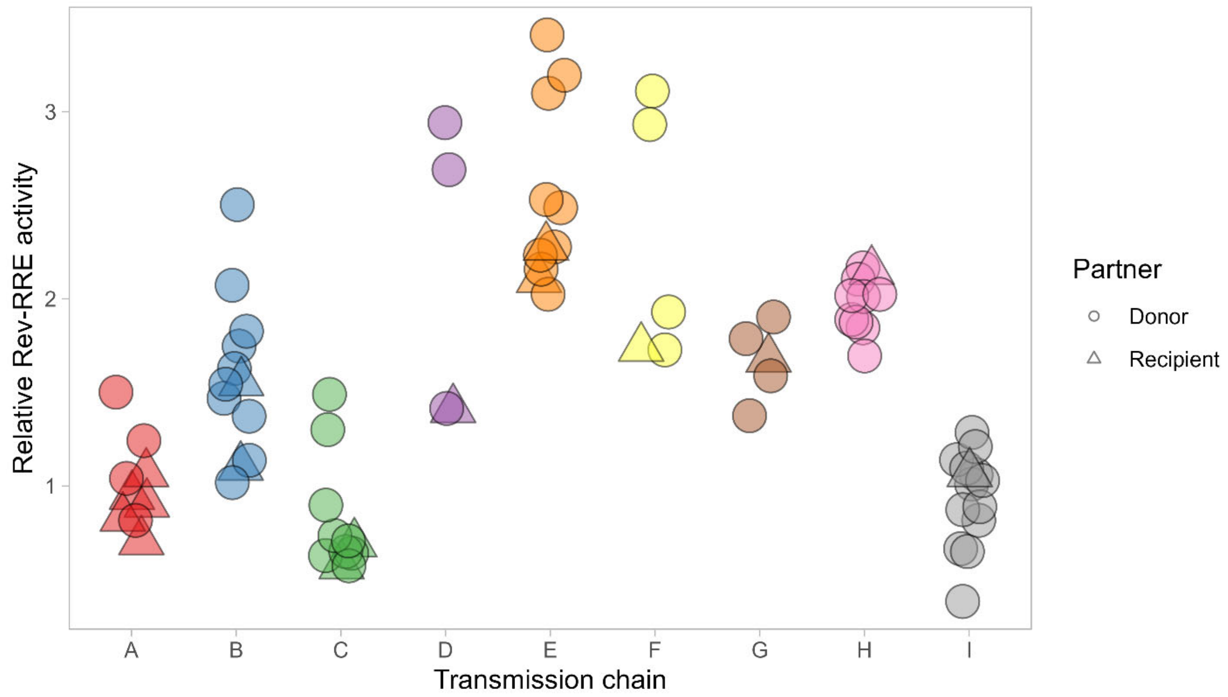


Figure S4. Rev-RRE functional activity of viral variants across transmission chains. Relative Rev-RRE functional activity of all pairs visualized in Figure 2 is again presented using a consistent y-axis for ease of comparison across the entire set. Rev-RRE pairs occurring within different transmission chains, A-I, are separated along the x axis and by color. Each symbol represents a unique Rev-RRE pair in a single individual. Relative activity is indicated on the y-axis in multiples of the functional activity of the NL4-3 Rev-RRE cognate pair. Rev-RRE sequences from donors or recipients are indicated by shape. Unlike in Figure 2, symbols are not sized based on the frequency of the Rev-RRE sequence within a quasispecies.
